# A generalized framework of AMOVA with any number of hierarchies and any level of ploidies

**DOI:** 10.1101/608117

**Authors:** Kang Huang, Yuli Li, Derek W. Dunn, Pei Zhang, Baoguo Li

## Abstract

The analysis of molecular variance (AMOVA) is a widely used statistical model in the studies of population genetics and molecular ecology. The classical framework of AMOVA only supports haploid and diploid data, in which the number of hierarchies ranges from two to four. In practice, natural populations can be classified into more hierarchies, and polyploidy is frequently observed in contemporary species. The ploidy level may even vary within the same species, even within the same individual. We generalized the framework of AMOVA such that it can be used for any number of hierarchies and any level of ploidy. Based on this framework, we present four methods to account for the multilocus genotypic and allelic phenotypic data. We use simulated datasets and an empirical dataset to evaluate the performance of our framework. We make freely available our methods in a software, POLYGENE, which is freely available at https://github.com/huangkang1987/.

## Introduction

The *analysis of molecular variance* (AMOVA) is a statistical model for the molecular variation in a single species. AMOVA was developed by Excoffier *et al.* (1992) based on the previous work of decomposing the total variance of gene frequencies into the variance components in different subdivision levels (Cockerham 1969; Cockerham 1973).

This statistical model was initially implemented for DNA haplotypes, but it can be applied to any marker datum, e.g., the codominant marker data and the dominant marker data (Peakall *et al.* 1995; Peakall and Smouse 2006). The classical framework of AMOVA supports haploids and diploids, and the number of hierarchies ranges from two to four (individual, population, group and total population) (Excoffier and Lischer 2010). In practice, natural populations can be classified into more than four hierarchies, and the ploidy level may vary within the same species or within the same individual.

In many species, physical or ecological barriers prevent random mating (Martin and Willis 2007). The resulting partial or total isolation of populations results in genetic differentiation due to the interacting processes of genetic drift, differential gene-flow and natural selection (Lande 1976). Because the factors restricting gene flow, such as geographical distance (Wright 1943), landscape features (e.g., mountain, river, desert) (Geffen *et al.* 2004; Chambers and Garant 2010; Lait and Burg 2013), ecological factors (e.g., salt concentration, climatic gradients) (Luppi *et al.* 2003; Yang *et al.* 2014) and behavioral differences (e.g., parental care) (Russell *et al.* 2004), are not all the same among populations, the gene flow between populations is unevenly distributed. For example, in humans, the intra-city gene-flow is higher than the intra-province, intra-nation and inter-nation gene flows. The population structure, in some situations, can be classified into multilevel hierarchies.

Polyploids represent a significant portion of plant species, with anywhere between 30% and 80% of angiosperms showing polyploidy (Burow *et al.* 2001) and most lineages showing the evidence of paleopolyploidy (Otto 2007). Due to their significant roles in molecular ecology, evolutionary biology and agriculture studies, polyploids have increasingly become the focus of theoretical and experimental research (Avni *et al.* 2017; Ling *et al.* 2018). There are two major problems in the population-genetics analysis of polyploids: genotyping ambiguity and double reduction.

For the *polymerase chain reaction* (PCR)-based markers, because the dosage of alleles cannot be determined by electrophoresis bands, the true genotype cannot be identified from the electrophoresis. This phenomenon is called *genotyping ambiguity*. For example, if an autotetraploid genotype *AAAB* has the same electrophoresis band type as the genotype *AABB*, then these two genotypes cannot be distinguished by electrophoresis.

In polyploids, double reduction occurs when a pair of sister chromatids is segregated into the same chromosome, which will cause the corresponding genotypic frequency deviating from *Hardy-Weinberg equilibrium* (HWE), where we assume that each allele will randomly appear within various genotypes. For autotetraploids, the rate α of double-reduction is assumed to be 0 under HWE, 1/7 under *random chromosome segregation* (RCS) (Haldane 1930), and 1/6 under *complete equational segregation* (CES) (Mather 1935). In the *partial equational segregation* (PES) model, the distance between the target locus and the centromere is incorporated into CES (Huang *et al.* 2019).

Some software for the polysomic inheritance model has been developed, e.g., POLYSAT (Clark and Jasieniuk 2011), SPAGEDI (Hardy and Vekemans 2002), POLYRELATEDNESS (Huang *et al.* 2014), GENODIVE (Meirmans and Tienderen 2004; Meirmans and Liu 2018), and STRUCTURE (Pritchard *et al.* 2000). However, some of them cannot solve the genotyping ambiguity, and all of them are supposed that the genotypic frequencies accord with HWE.

In this paper, we generalize the framework of AMOVA such that any number of hierarchies and any level of ploidy are allowed. Four methods are developed to account for multilocus genotypic and phenotypic data, including three method-of-moment methods and one maximum-likelihood method. Our model has been implemented in a software named POLYGENE, and it is freely available at https://github.com/huangkang1987/. POLYGENE is designed for genotypic or phenotypic datasets, which only supports homoploids to include more population-genetics analyses (e.g., phenotypic/genotypic distribution test).

## Theory and modeling

There are three purposes of typical AMOVA: (i) estimate the variance components in different subdivision levels; (ii) measure the population differentiation with *F*-statistics (*F*_*IS*_, *F*_*IT*_, *F*_*ST*_, etc.); (iii) test the significance of differentiation. In the following sections, we will briefly describe the general procedures of classic framework of AMOVA, then extend them to the generalized situation.

### Classic framework

The procedures of AMOVA are as follows: (i) calculate the genetic distance between two alleles or two haplotypes; (ii) calculate the *sum of squares* (SS), the degree of freedom and the *mean square* (MS) in each source of variation; (iii) solve variance components; (iv) calculate *F*-statistics; (v) perform permutation tests.

The collection consisting of some populations is called a *group*, denoted by *g*. We stipulate that each population can only belong to one group, and the union of all groups is the total population. Because an allele or a haplotype (for simplicity, we use ‘allele’ to refer them hereafter) is usually neither a discrete nor a continuous random variable (except the allele size in microsatellites), the SS cannot be calculated by the equation SS = ∑_*i*_(*X*_*i*_ − *X̅*)^2^. Using the genetic distance between any two alleles as a proxy, an alternative method can be used to calculate the SS, whose formula for a group of *n* allele copies is as follows:

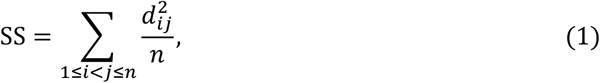

where *d*_*ij*_ is the genetic distance between the *i*^th^ and the *j*^th^ alleles. Such genetic distance is one of the following distances: nucleotide difference distance for DNA sequences (DNA sequence, Excoffier *et al.* 1992), Euclidean distance for dominant markers (dominant marke, Peakall *et al.* 1995), *infinity allele model* (IAM) distance for codominant marker (codominant marker, Cockerham 1973) and *stepwise mutation model* distance (SMM) for microsatellites (microsatellite, Slatkin 1995).

In variance decomposition, the genetic variance is decomposed as two to four hierarchies, including 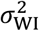 (within individuals), 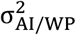 (among individuals within populations), 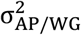 (among populations within groups) and 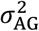 (among groups), where 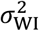 and/or 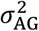 are sometimes ignored. Using all of the four hierarchies as an example, the layout of AMOVA is shown in Table 1.

**Table 1.**
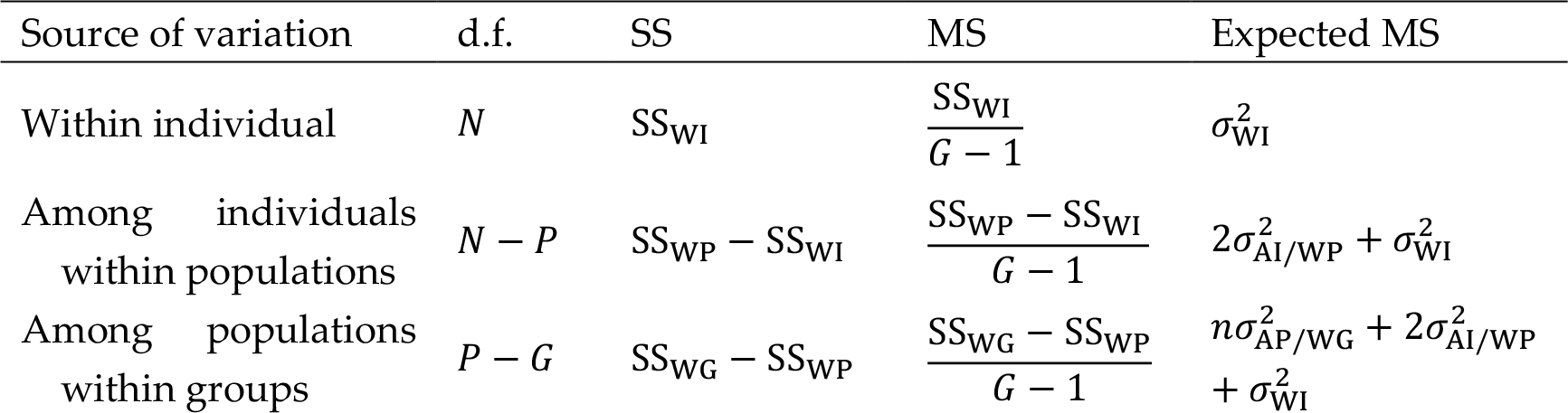
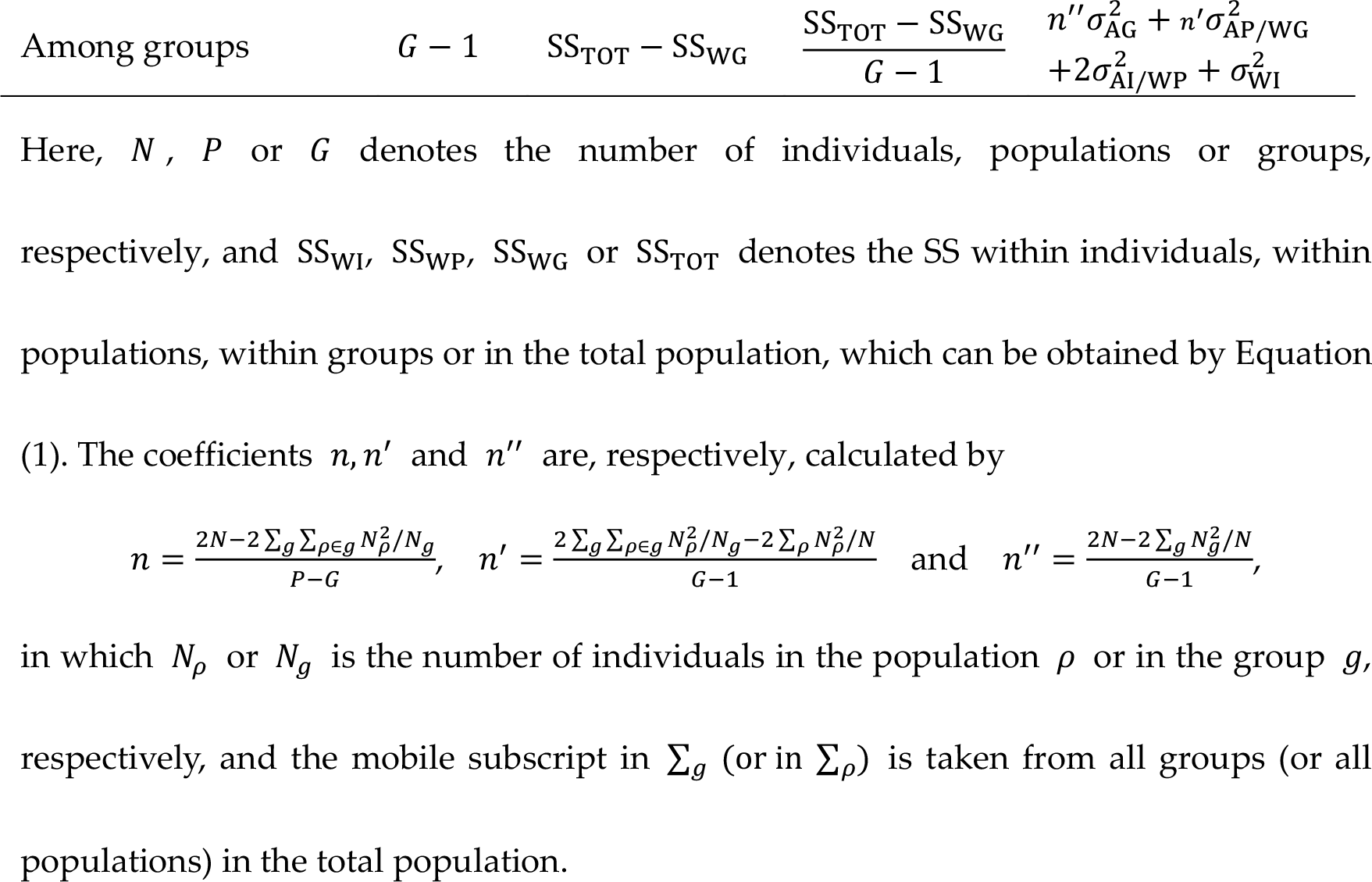
The layout of AMOVA. The total SS is decomposed into the SS in different sources of variation. Each expected MS is expressed here as a function of variance components.

| Source of variation | d.f. | SS | MS | Expected MS |
| --- | --- | --- | --- | --- |
| Within individual | $N$ | $SS_{WI}$ | $\frac{SS_{WI}}{G - 1}$ | $\sigma_{WI}^2$ |
| Among individuals<br>within populations | $N - P$ | $SS_{WP} - SS_{WI}$ | $\frac{SS_{WP} - SS_{WI}}{G - 1}$ | $2\sigma_{AI/WP}^2 + \sigma_{WI}^2$ |
| Among populations<br>within groups | $P - G$ | $SS_{WG} - SS_{WP}$ | $\frac{SS_{WG} - SS_{WP}}{G - 1}$ | $n\sigma_{AP/WG}^2 + 2\sigma_{AI/WP}^2 + \sigma_{WI}^2$ |
| Among groups | $G - 1$ | $SS_{TOT} - SS_{WG}$ | $\frac{SS_{TOT} - SS_{WG}}{G - 1}$ | $\frac{n''\sigma_{AG}^2 + n'\sigma_{AP/WG}^2}{+2\sigma_{AI/WP}^2 + \sigma_{WI}^2}$ |

Here, $N$ , $P$ or $G$ denotes the number of individuals, populations or groups, respectively, and $SS_{WI}$ , $SS_{WP}$ , $SS_{WG}$ or $SS_{TOT}$ denotes the SS within individuals, within populations, within groups or in the total population, which can be obtained by Equation (1). The coefficients $n, n'$ and $n''$ are, respectively, calculated by
in which $N_{\rho}$ or $N_g$ is the number of individuals in the population $\rho$ or in the group $g$ , respectively, and the mobile subscript in $\sum_g$ (or in $\sum_{\rho}$ ) is taken from all groups (or all populations) in the total population.

By equating the expected MS with the observed MS, the unbiased estimates of variance components can be solved. After that, the *F*-statistics can be calculated by the following formulas:

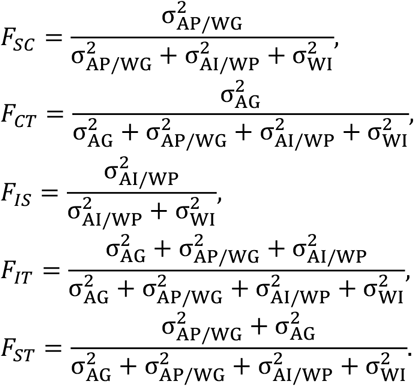

A permutation test is used to test the significance of differences. The null hypothesis is that there is no differentiation among individuals, populations or groups, and the observed differences are due to the random sampling. This statement is equivalent to the variance 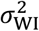 occupying 100% of the total variance, and thus the variances 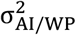, 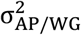 and 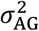 together with various *F*-statistics are all zero.

In each permutation, the allele copies are randomly permuted in the total population to generate a new dataset. The variance components and the *F*-statistics are calculated for each permuted dataset to obtain their distributions under null hypothesis. The probability that each permuted variance component or each *F*-statistic is greater than the original value is used as a single-tailed *P*-value.

### Generalized framework

Here, we present a generalized framework to decompose the genetic variance. In our framework, we do not need to use the degrees of freedom or the MS. Instead, we directly use the variance components to express the expected SS of various hierarchies, whose expressions are as follows:

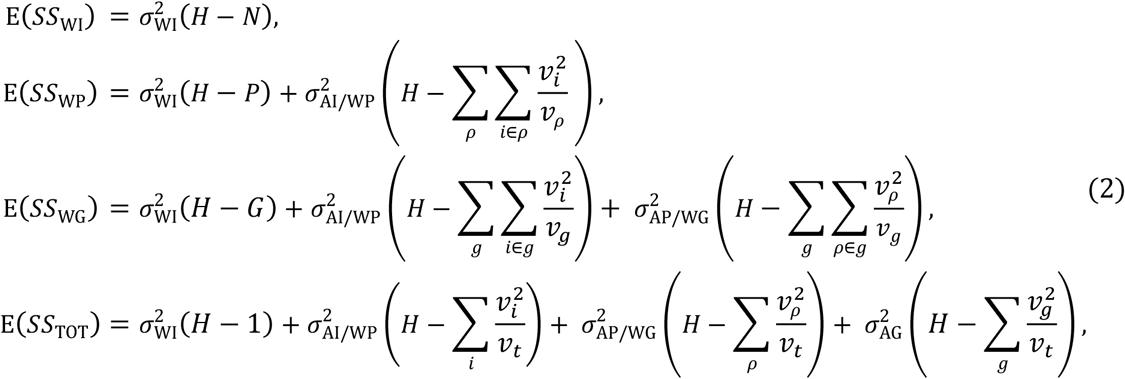

where *H* and *ν*_*t*_ are, respectively, the total number of haplotypes and alleles, *ν*_*i*_ (*ν*_*ρ*_ or *ν*_*g*_) is the number of haplotypes/alleles within the individual *i* (the population *ρ* or the group *g*), and the mobile subscript in ∑_*ρ*_ (∑_*g*_ or ∑_*i*_) is taken from all populations (all groups or all individuals). The estimates of variance components are identical to those in the classic framework. The step-by-step detailed derivations for these formulas in Equation (2) are provided in the Supplementary materials. Compared with previous frameworks (Table 1), Equation (2) is more regular, making it possible to be generalized.

According to the concept that each member in a hierarchy is a ‘vessel’ of genes, we can use the vessels *V*_0_, *V*_1_, *V*_2_, *V*_3_, *V*_4_ and the corresponding expected SS_*i*_ (*i* = 1, 2, 3, 4) to describe each formula in Equation (2). In other words, we can use these vessels to rewrite Equation (2):

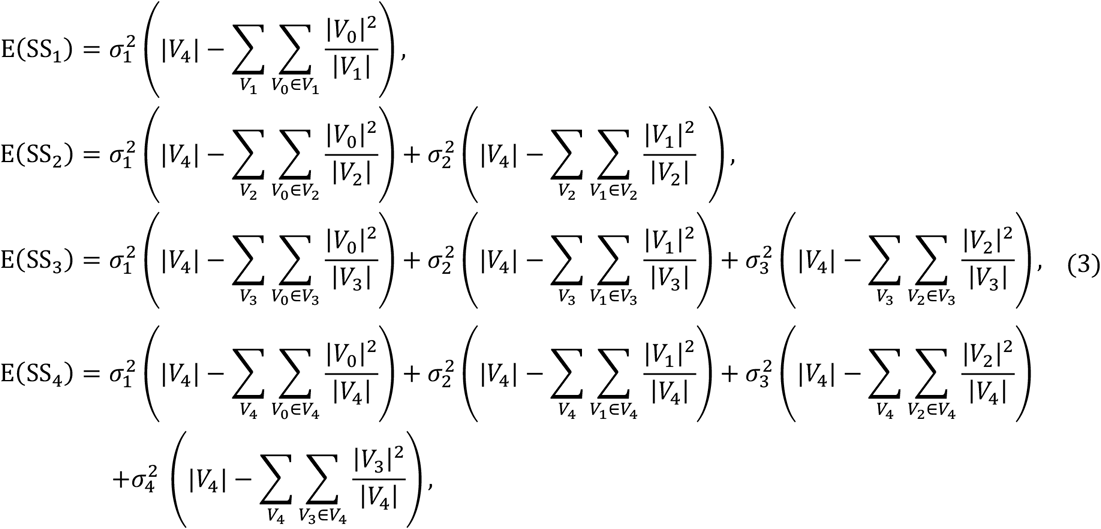

where *V*_*i*_ is a vessel at the level *i*, |*V*_*i*_| is the number of allele copies in *V*_*i*_, SS_*i*_ is the SS within all vessels at the level *i*, 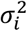 is the variance component among all *V*_*i*−1_ within *V*_*i*_, and the mobile subscript in 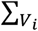 is taken from all vessels at the level *i*. The subscript *i* ranges from 0 to 4, and the corresponding vessels represent, in turn, alleles, individuals, populations, groups and the total population. Equation (3) is in apple-pie order, which can be expressed as the forms of summation signs:

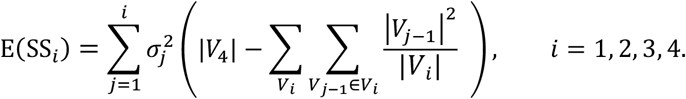

If the hierarchy of individuals is ignored, then the vessel *V*_1_ (*V*_2_ or *V*_3_) will represent a population (a group or the total population). In this situation, Equation (3) becomes

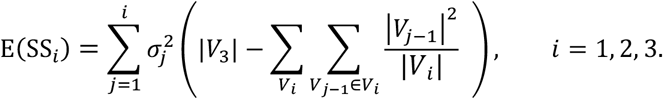

Generally, if there are *M* + 1 kinds of vessels at the levels ranging from 0 to *M*, then Equation (3) can be generalized as follows:

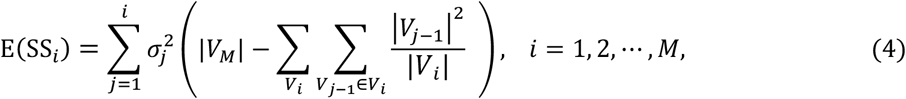

where *V*_*M*_ denotes the vessel of highest hierarchy (i.e., the total population). Equation (4) is the ultimate generalized form of AMOVA, which is extremely simple and can be applied to any number of hierarchies and any level of ploidy. We can also use matrices to express Equation (4):

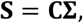

where **S** = [E(*SS*_1_), E(*SS*_2_),…, E(*SS*_*M*_)]^*T*^, 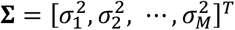 and the coefficient matrix **C** is lower-triangular with type *M* × *M*, whose *ij*^th^ element is

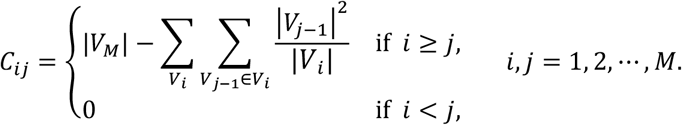

Then, a method-of-moment estimation of variance components can be given by 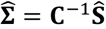, and the *F*-statistics can be solved by

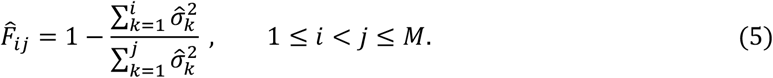

### Method-of-moment methods

For convenience, we call a method-of-moment estimator a *moment method*. In practice, the multilocus data are used to increase the accuracy of estimation. Based on the moment estimator described above, we develop three methods (called the *homoploid, anisoploid* and *weighting genotypic methods*) to account for the multilocus genotypic or phenotypic data.

The **homoploid method** is only applicable to homoploids. In this method, all loci are treated as one dummy locus, and the dummy haplotypes are extracted from phenotypes. Meanwhile, the genetic distance between any two dummy haplotypes is calculated, and these dummy haplotypes are permuted to test the significance of differentiation. In diploids, this method is used in GENALEX (Peakall and Smouse 2006).

To solve the genotyping ambiguity, we will use the posterior probabilities to weight the possible genotypes hidden behind a phenotype. The multiset consisting of alleles within an individual and at a locus is defined as a *genotype*, denoted by 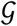, and the set obtained by deleting the duplicated alleles in 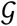 is defined as the *phenotype* determined by 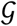, denoted by 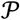. In our previous paper (Huang *et al.* 2019), the genotypic frequency 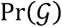 and the phenotypic frequency 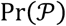 under a double-reduction model (HWE, RCS, CES or PES) were calculated. On this basis, we are able to calculate the posterior probability 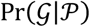 of a genotype 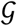 determining 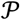, whose formula is as follows:

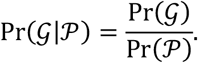

After that, the probability Pr(*A*_*hl*_ = *A*_*lj*_) (or *p*_*hlj*_ for short) can be calculated by

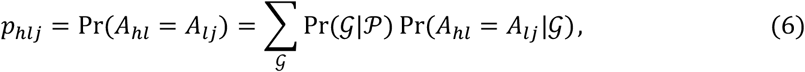

where *A*_*hl*_ is the allele in the *h*^th^ dummy haplotype and at the *l*^th^ locus, *A*_*lj*_ is the *j*^th^ allele at the *l*^th^ locus, and 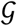 is taken from all possible genotypes determining 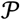.

In the homoploid method, because all loci are treated as one dummy locus, the square of genetic distance *d*_*hh*′_ between the *h*^th^ and the 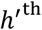 haplotype is the sum of squares of the distances in the form 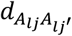 over all *L* loci, namely

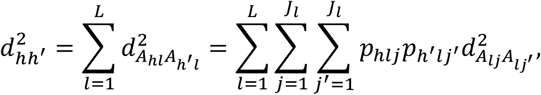

where 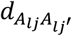 is the distance between the *j*^th^ and the 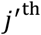 allele at the *l*^th^ locus, and *J*_*l*_ is the number of alleles at the *l*^th^ locus. For an allele with missing data, its frequency refers the frequency in the corresponding population. In this method, because there is only one dummy locus, no additional weighting procedure is required for multilocus data.

The **anisoploid method** can be applied for both homoploids and anisoploids. In this method, the dummy alleles are extracted at each locus, and the missing data are ignored. Meanwhile, the genetic distance between two dummy alleles needs to be calculated locus by locus, and the dummy alleles are randomly permuted locus by locus during the permutation test.

For this method, the probability Pr(*A*_*h*_ = *A*_*j*_) (or *p*_*hj*_ for short) in a phenotype at a target locus can also be expressed by Equation (6). Then the square of genetic distance 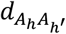 between two allele copies *A*_*h*_ and *A*_*h*′_ at this target locus is given by

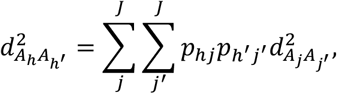

where *J* is the number of alleles at this target locus.

The untyped individuals (populations or groups) due to missing data should be directly skipped to avoid a denominator of zero. The global variance components for multilocus data can be solved by using the formula **S** = **CΣ**. There are two solving strategies: (i) find the sum of the whole **S** and the sum of the whole **C** over all loci, and then solve the global variance components, denoted by 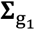; (ii) solve **Σ** for each locus, and then find the sum of whole **Σ** over all loci, denoted by 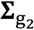. Generally, 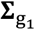 and 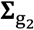 are different, but they are approximately proportional to each other.

We adopt the first strategy because the global SS, the d.f. and the MS can also be obtained. This strategy has the same output style as the classic framework.

In the **weighting genotypic method**, no dummy haplotypes are extracted. Instead, for any hierarchy *i*, the SS_*i*_ for each genotype hidden behind a phenotype is calculated, and then all sums of squares in the hierarchy *i* are weighted according to the corresponding posterior probabilities. We also find the sum of those SS_*i*_ over all loci in this method. For each locus, the SS_*i*_ is calculated by

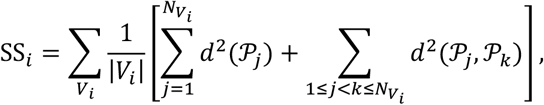

where SS_1_ = SS_WI_ (i.e., when *i* = 1), *V*_*i*_ is taken from all vessels in the hierarchy 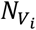 is the number of phenotypes determined by the individuals within *V*_*i*_, and at this locus, 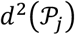(or 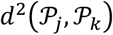) is the weighted sum of squares of the distances within the phenotype 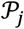 (or between the phenotypes 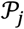 and 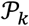), which can be calculated by the following formulas:

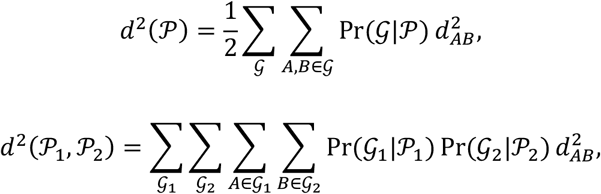

where 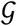 (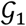 or 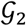) is taken from all candidate genotypes determining 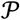 (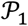 or 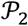), and *d*_*AB*_ is the distance between the alleles *A* and *B*.

### Maximum-likelihood method

We will develop a maximum-likelihood estimator to estimate the *F*-statistics and solve the variance components. For convenience, we call this method the *likelihood method*. In this method, a reversed procedure is used, such that the *F*-statistics are first estimated, and next the variance components and other statistics are solved.

To derive the expression of the likelihood for individuals, we first model some equations. A random distribution can be used to simulate the differentiation among individuals within a vessel *V*_*i*_. We will choose some Dirichlet distribution for each hierarchy. That is because the standardized variance of each allele frequency accords with the corresponding *F*-statistic in that distribution. Therefore, no additional weighting procedure for the variance components is required.

Given a vessel *V*_*i*_ (2 ≤ *i* ≤ *M*), the allele frequencies *p*_11_, *p*_12_,…, *p*_1*J*_ within an individual (i.e., within one of those *V*_1_) are drawn from the Dirichlet distribution 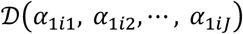, where *J* is the number of alleles within this individual, and

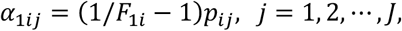

in which *F*_1*i*_ is the *F*-statistic among all individuals within *V*_*i*_, and *p*_*ij*_ is the frequency of *j*^th^ allele in *V*_*i*_. Then, the expectation and the variance of *j*^th^ allele frequency *p*_1*j*_ as a random variable are, respectively, *p*_*ij*_ and *F*_1*i*_*p*_*ij*_(1 − *p*_*ij*_), and the standardized variance of *p*_*ij*_ as a random variable is exactly *F*_*i*,*i*+1_, which is identical to Wright’s definition of *F*-statistics.

For simplicity, we let **p**_*i*_ be the vector consisting of the frequencies of all alleles in *V*_*i*_, i.e.

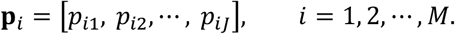

Then, for each *i* with 2 ≤ *i* ≤ *M*, the probability density function of **p**_1_ is as follows:

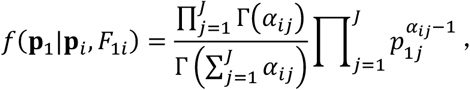

where Γ(⋅) is the gamma function, and α_*ij*_ = 1/F_*ij*_ − 1. Assume that the alleles within *V*_1_ are independently drawn according to the frequencies in **p**_1_. Then, the allele copy numbers of *V*_1_ obey a multinomial distribution, and so the frequency 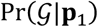 of a genotype 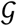 conditional on **p**_1_ is

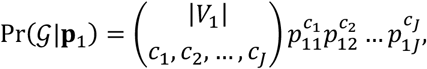

where *c*_*j*_ is the number of the *j*^th^ allele copies in 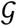, *j*= 1, 2,…, *J*.

Now, the frequency 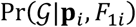 of 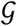 conditional on both **p**_*i*_ and *F*_1*i*_ can be obtained from the weighted average of 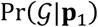 with *f*(**p**_1_|**p**_*i*_, *F*_1*i*_)d**p**_1_ as the weight, that is,

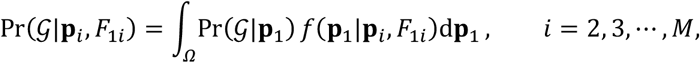

where the integral domain *Ω* can be expressed as

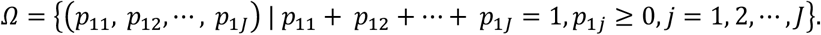

The integral can be converted into the following repeated integral with the multiplicity *J* − 1:

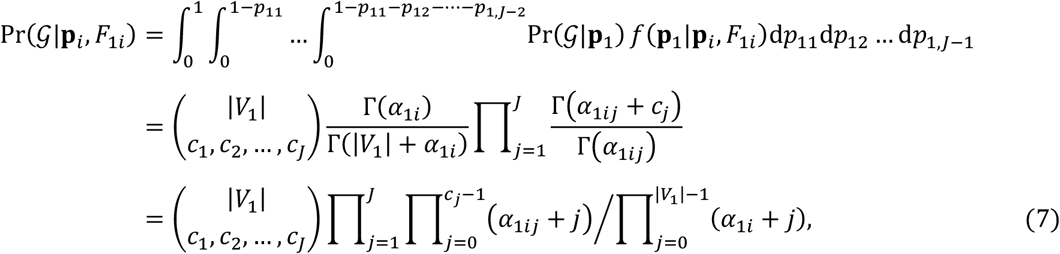

where *α*_1*i*_ = 1/*F*_1*i*_ − 1. Importantly, *α*_1*i*_ → +∞ if *F*_1*i*_ → 0^+^, thus the variance 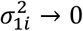 if *F*_1_*i* → 0^+^. Since **p**_*i*_ is unavailable, the estimate 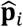 is used as **p**_*i*_ in our calculation.

The frequency 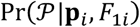 of a phenotype 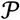 conditional on both **p**_*i*_ and *F*_1*i*_ is the sum of frequencies in the form 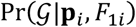, where 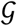 is taken from all candidate genotypes determining 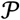, in other words,

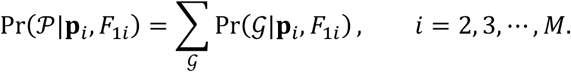

Now, the global likelihood for individuals at a hierarchy *i* can be obtained, which is the product of frequencies in the form 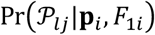 over all individuals and at all loci, symbolically

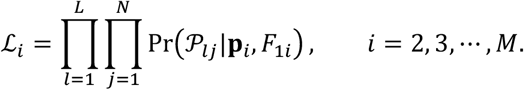

Because the allele frequencies are already estimated, a downhill simplex algorithm (Nelder and Mead 1965) can be used to find the optimal *F*_1*i*_ under the IAM model. After that, the variance components can be solved from the *F*-statistics with an additional constraint as follows:

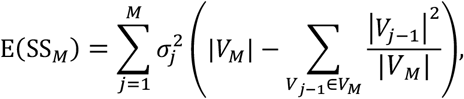

where SS_*M*_ can be obtained from the allele frequencies of the total population under the IAM model, that is,

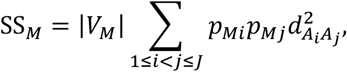

where 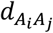 is the IAM distance between the alleles *A*_*i*_ and *A*_*j*_.

### Differentiation test

In the homoploid/anisoploid method, the dummy haplotypes/alleles are extracted. Then, the differentiation test can be performed by permuting the dummy haplotypes/alleles. However, for the weighting genotypic and the likelihood methods, this cannot be done because there are neither dummy haplotypes nor dummy alleles being extracted in these two methods.

To solve this problem, we develop an alternative method to test the differences. In this method, the datasets of the same structure as the original datasets are randomly generated, where ‘the same structure’ means that there are the same individuals, populations and groups as well as the same missing data. More specifically, the genotypes of each individual are generated conditional on **p**_*M*_ and *F*_1*M*_ according to Equation (7) under the null hypothesis that there are no differences (i.e., *F*_1*M*_ → 0^+^). Moreover, the phenotypes can be obtained by removing the duplicated alleles in the generated genotypes.

For each generated dataset, the variance components and the *F*-statistics are estimated by the same procedures as above to obtain their empirical distributions. Similarly, the probability that each permuted variance component or each *F*-statistic is greater than the original value is used as a single-tailed *P*-value.

The authors affirm that all data necessary for confirming the conclusions of the article are present within the article, figures, and tables.

## Evaluations

### Simulated data

A Monte-Carlo simulation is used to assess the accuracy of the four methods mentioned above (three moment methods and one likelihood method) under different conditions: ploidy level, number of hierarchies and population differentiations. We choose three types of hierarchies: *M* = 3, 4 or 5. If we denote by *n*_*i*_ the number of vessels in the form *V*_*i*−1_ in *V*_*i*_, then the ploidy level *n*_1_ of each individual (i.e., the number of allele copies in each individual at a locus) is set as *n*_1_ = 2, 4 or 6, and the number *n*_2_ of individuals sampled in each population ranges from 5 to 50 at intervals of 5. For those higher-hierarchy vessels, we set *n*_3_ = 4 and *n*_4_ = *n*_5_ = 2. In the following discussion, we will use *N* to replace the symbol *n*_2_. Meanwhile, we set the number of loci per population as 10 and set the number of alleles per locus *J* = 6. We simulate these three types of hierarchies at each of the three ploidy levels in turn.

For the total population (i.e., *V*_*M*_), the allele frequencies *p*_*M*1_, *p*_*M*2_,…, *p*_*MJ*_ (*M* = 3, 4 or 5, *J* = 6) are randomly drawn from the Dirichlet distribution 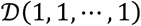 with all concentration parameters being equal to one. The *F*-statistic *F*_*i*,*i*+1_ among all *V*_*i*_ within *V*_*i*+1_ is set as 0.05. To simulate the differentiation, the allele frequencies in **p**_*i*_ for each *V*_*i*_ are independently generated according to both **p**_*i*+1_ and *F*_*i*,*i*+1_. More specifically, *p*_*i1*_, *p*_*i2*_,…, *p*_*iJ*_ are randomly drawn from the Dirichlet distribution 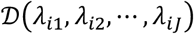, *i* = 1, 2,…, *M* − 1, where

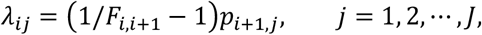

in which *p*_*i*+1,*j*_ is the frequency of *j*^th^ allele in the upper vessel *V*_*i*+1_. Obviously, each *λ*_*ij*_ is proportional to *p*_*i*+1,*j*_.

The alleles in each individual are randomly drawn according to the allele frequencies *p*_11_, *p*_12_,…, *p*_1*J*_ for this individual (i.e., one of the vessels in the form *V*_1_). For polyploids, the duplicated alleles within a genotype 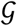 will be removed to convert 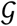 into a phenotype 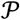. The genotype 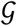 or the phenotype 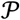 is randomly set as ∅ at a probability of 0.05 to simulate the negative amplification.

For any combination of simulation parameters, 5,000 datasets are generated, and then the AMOVA for every generated dataset is performed by using each of these four methods. The allele frequencies for each population at each locus are independently estimated by using the double-reduction model under RCS with inbreeding. An expectation-maximization algorithm modified from Kalinowski and Taper (2006) is used to estimate the frequencies of alleles. In this algorithm, the initial value of each allele frequency at a target locus is assigned as 1/*J*, and then each frequency is iteratively updated until the sequence consisting of those updated values is convergent. The updated frequency 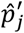 is calculated by

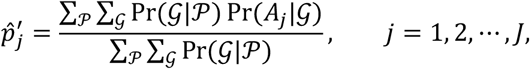

where 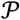 is taken from all phenotypes at this target locus, 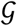 is taken from all possible genotypes determining 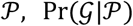 is the posterior probability of 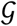 determining 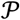, and 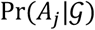 is the frequency of the *j*^th^ allele in 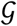 and at this target locus.

Because the dimension of variance components depends on the allele frequencies and the number of loci, to estimate the *F*-statistics, what we truly need is the proportion 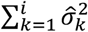 to 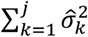 according Equation (5). Therefore, we use the bias and the RMSE of the *F*-statistics to evaluate the accuracy of estimates of *F*-statistics, where RMSE is the abbreviation of *root-mean-square error*.

### Simulated results

The bias and the RMSE of 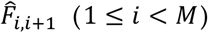 for diploids and under different conditions are shown in Figures 1 and 2, respectively. It can be found from Figure 1 that each bias is generally reduced as the sample size *N* increases, and its variation trend does not change as the number *M* of hierarchies increases. The bias of 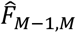 is smaller than that of 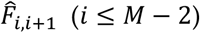. For the anisoploid and the weighting genotypic methods, the estimates of *F*-statistics are unbiased. However, for the homoploid method, they are slightly biased due to the weighting for missing data. For the likelihood method, the bias is largest, reaching 0.05, but it drops to below 0.015 at *N* = 50. As *M* increases, the bias of 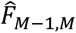 is within the range of the other *F*-statistics.

**Figure 1.**
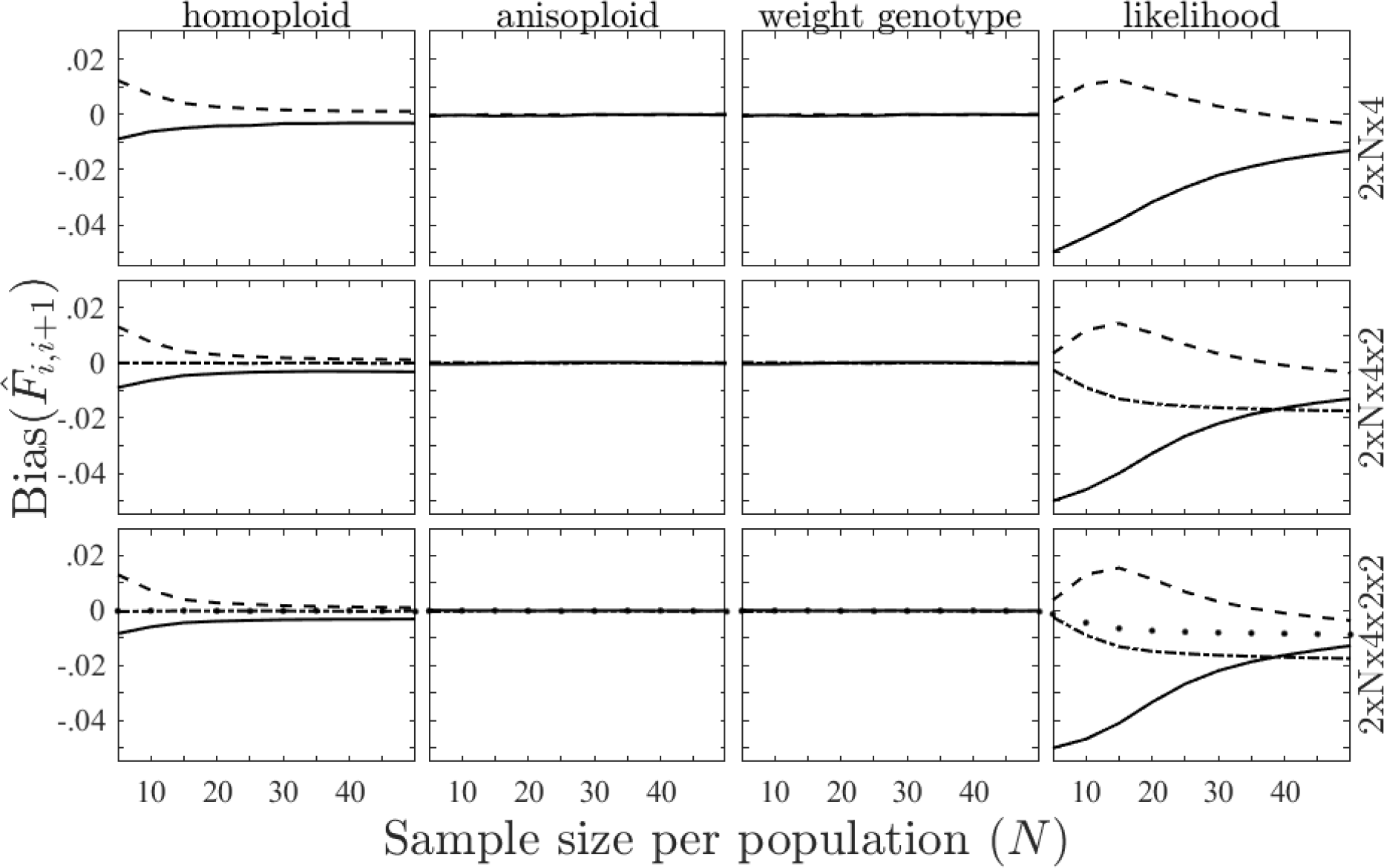
The bias of estimated *F*-statistic 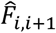 for diploids as a function of sample size *N* sampled from each population under different conditions (1 ≤ *i* < *M*). For each of the four methods listed at the top, the results are shown in the column where this method is located. Each row shows the results of a structure, where the structure ‘2×N×4×2’ means that *M* = 4, and there are two groups, each group containing four populations and each population consisting of *N* diploids. The meanings of other two structures can be analogized. Each solid, dashed, dash-dotted or dotted line denotes the bias of 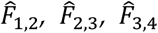 or 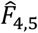, respectively, corresponding to the value of *N*.

**Figure 2.**
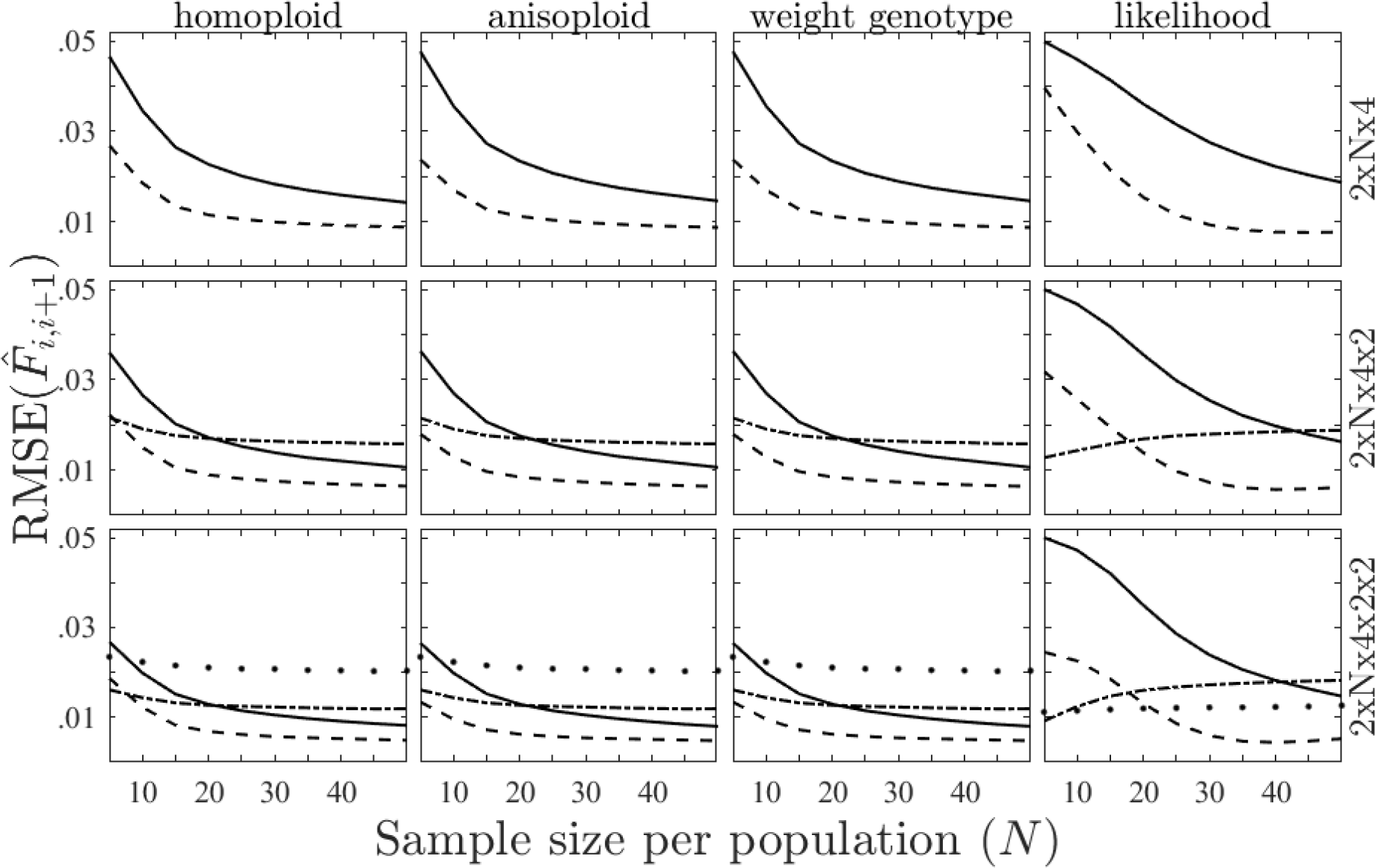
The RMSE of estimated *F*-statistic 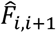 for diploids and under different conditions (1 ≤ *i* < *M*). The meanings of columns and rows are as indicated in Figure 1. Each solid, dashed, dash-dotted or dotted line denotes the RMSE of 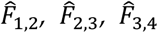 or 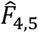, respectively, corresponding to the value of *N*.

It can be seen from Figure 2 that for the three moment methods, the RMSEs of 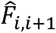 are similar: if *M* is lower (e.g., *M* = 3), the variance decreases quickly as *N* increases. However, if *M* is larger, the RMSE of 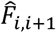 is less sensitive to the changes in *N*, and the RMSE of 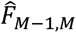 becomes more and more inaccurate as *M* increases. In contrast, for the likelihood method, the RMSE of 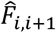 is less affected by *M*, and the RMSE of 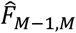 lies among those of 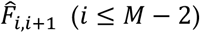.

The bias and the RMSE of 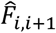 for tetraploids and under different conditions are shown in Figures 3 and 4, respectively. It can be seen from Figure 3 that for the three moment methods, the estimates of *F*-statistics become biased for the polyploid phenotypic data. The bias of x*F*_1,2_ is largest, reaching −0.01 at *N* = 50. For the weighting genotypic method, the estimates of *F*-statistics are also biased, but their biases drop to 0.003 at *N* = 50. For the likelihood method, the bias is larger than that of the weighting genotypic method, reaching 0.02 at *N* = 50. As *M* increases, the bias of 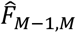 is also around those of the other *F*-statistics.

**Figure 3.**
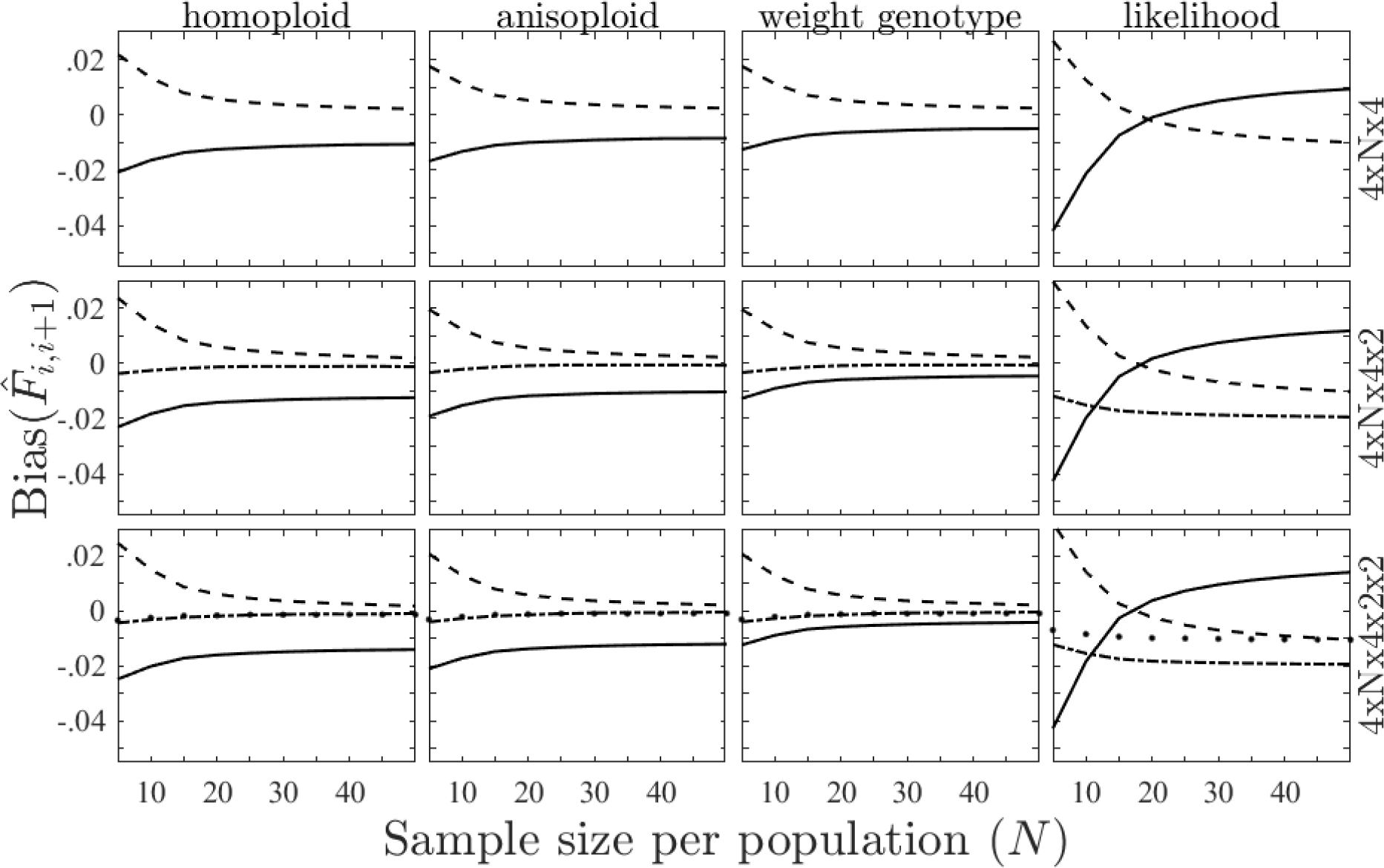
The bias of estimated *F*-statistic 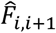 for tetraploids and under different conditions. The meanings of columns, rows and lines are as indicated in Figure 1.

**Figure 4.**
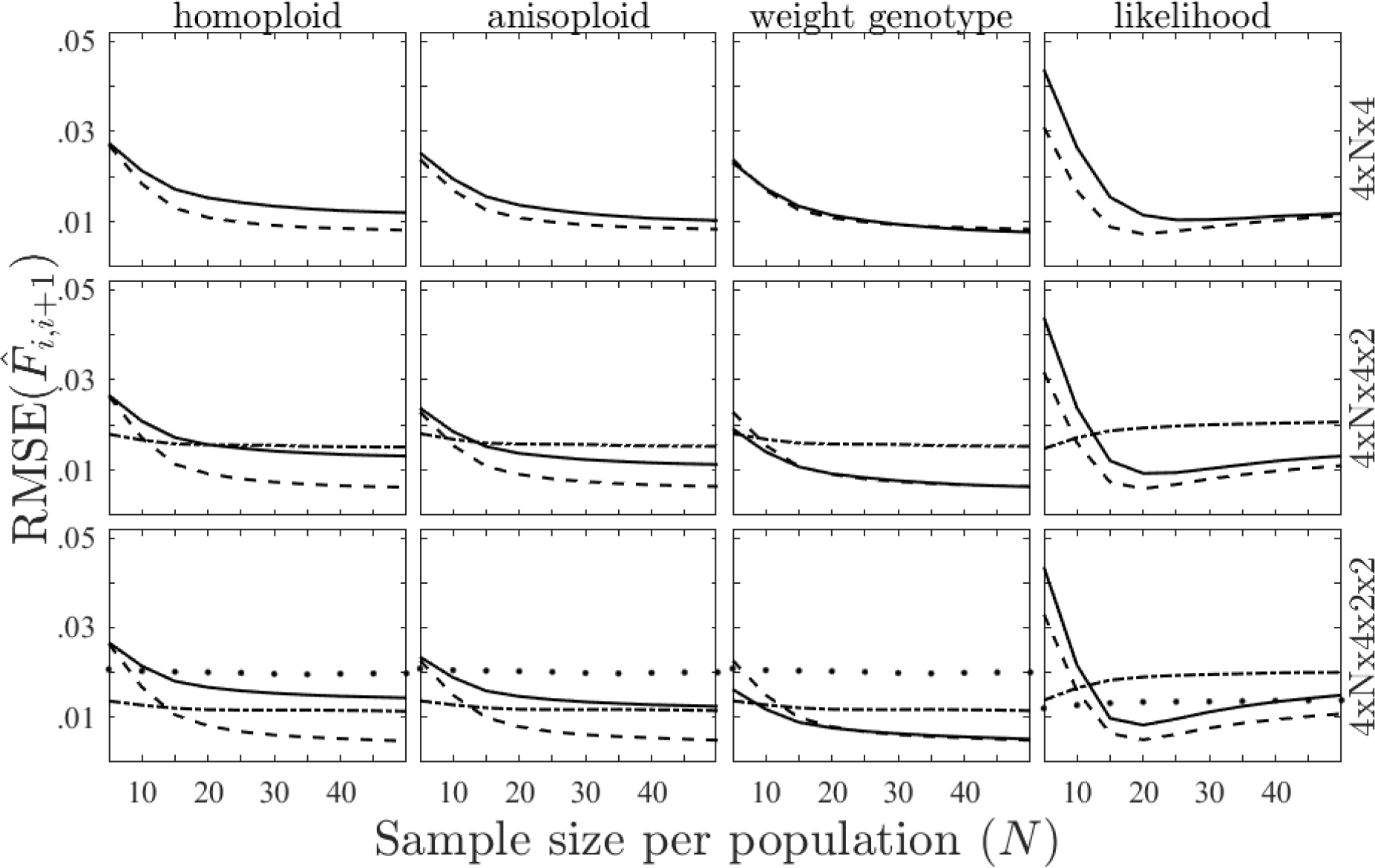
The RMSE of estimated *F*-statistic 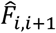 for tetraploids and under different conditions. The meanings of columns, rows and lines are as indicated in Figure 2.

Compared with the situation of diploids, the RMSE in Figure 4 is reduced in scale, while the patterns are similar to those in diploids. For the weighting genotypic method, the RMSE of 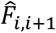 becomes less sensitive to *N* as *M* increases, and the RMSE of 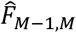 is largest. For the likelihood method, the sensitivity of RMSE of 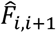 does not vary significantly as *M* increases.

### Empirical data

We will use the human dataset of Pemberton *et al.* (2013) to evaluate our generalized framework of AMOVA. This dataset consists of 5795 individuals sampled from 267 worldwide populations (e.g., ethnic groups). These populations are genotyped at 645 autosomal microsatellite loci. The average genotyping rate is 97.02%. In this dataset, the notion of groups needs to be divided into two levels, called *groups I* and *groups II*, to generate a nested structure with five levels (individual, population, group I, group II, total population).

The collection consisting of several populations in some countries or areas is defined as a *group I*, and the collection consisting of several populations in some region (e.g., East Asia or Middle East) is defined as a *group II*. For example, the populations of all Chinese nations are assigned to East Asia, whereas the population of the Uygur ethnic group is originally in Central South Asia. We still stipulate that each population (or each group I) can only belong to one group I (or one group II), and the union of all groups with the same level is the total population.

### Empirical results

Because Pemberton *et al.* (2013) dataset is genotypic and because 2.98% of genotypes are missing, the weighting genotypic method is equivalent to the anisoploid method, and the homoploid method is biased. We only use the anisoploid and the likelihood methods for this dataset, whose results are shown in Table 2. Moreover, the results of the corresponding *F*-statistics are shown in Table 3.

**Table 2.** The degrees of freedom, sum of squares (SS), mean squares (MS), estimated variance components (Var) and the variance percentage of (microsatellite, Slatkin 1995) dataset

| Method | Source | d.f. | SS | MS | Var | % | P | Perm. Mean | Perm. Var |
| --- | --- | --- | --- | --- | --- | --- | --- | --- | --- |
| Anisoploid | WI | 3626266 | 1293746.5 | 0.357 | 0.357 | 92.68 | 1.000 | 0.381 | 0.000 |
|  | AI/WP | 3454743 | 1262341.6 | 0.365 | 0.004 | 1.12 | 0.000 | 0.000 | 0.000 |
|  | AP/WC1 | 133491 | 87578.6 | 0.656 | 0.007 | 1.82 | 0.000 | 0.000 | 0.000 |
|  | AC1/WC2 | 32227 | 36005.9 | 1.117 | 0.003 | 0.77 | 0.000 | 0.000 | 0.000 |
|  | AC2 | 5160 | 86911.1 | 16.843 | 0.014 | 3.62 | 0.000 | 0.000 | 0.000 |
|  | Total | 7251887 | 2766583.6 | 0.381 | 0.385 | 100.00 | - | - | - |
| Likelihood | WI | 3626266 | 1310274.0 | 0.361 | 0.361 | 94.00 | 1.000 | 0.381 | 0.000 |
|  | AI/WP | 3454743 | 1248297.8 | 0.361 | 0.000 | 0.00 | 0.624 | 0.000 | 0.000 |
|  | AP/WC1 | 133491 | 88346.1 | 0.662 | 0.007 | 1.88 | 0.000 | 0.000 | 0.000 |
|  | AC1/WC2 | 32227 | 45688.1 | 1.418 | 0.005 | 1.26 | 0.000 | 0.000 | 0.000 |
|  | AC2 | 5160 | 73977.7 | 14.337 | 0.011 | 2.86 | 0.000 | 0.000 | 0.000 |
|  | Total | 7251887 | 2766583.6 | 0.381 | 0.384 | 100.00 | - | - | - |

\*WI: within individuals; AI/WP: among individuals within populations; AP/WC1: among populations within groups I; AC1/WC2: among groups I within groups II; AC2: among groups II.

**Table 3.** The value, significance, permuted mean and permuted variance of *F*-statistics

| $F$-statistics | Anisoploid | | | | likelihood | | | |
| --- | --- | --- | --- | --- | --- | --- | --- | --- |
| | Value | $P$ | Mean<br>$\times 10^{-6}$ | Var<br>$\times 10^{-8}$ | Value | $P$ | Mean<br>$\times 10^{-4}$ | Var<br>$\times 10^{-6}$ |
| $F_{IS}$ | 0.0119 | <0.0001 | 0.0425 | 7.3783 | 0.0000 | 0.1649 | 0.0000 | 0.0000 |
| $F_{IC1}$ | 0.0307 | <0.0001 | 1.1174 | 7.2052 | 0.0196 | <0.0001 | 1.7837 | 0.3049 |
| $F_{IC2}$ | 0.0384 | <0.0001 | 1.2765 | 7.1643 | 0.0323 | <0.0001 | 3.9708 | 0.5911 |
| $F_{IT}$ | 0.0732 | <0.0001 | 1.3279 | 7.1507 | 0.0600 | <0.0001 | 6.1550 | 1.0658 |
| $F_{SC1}$ | 0.0190 | <0.0001 | 1.0729 | 0.2298 | 0.0196 | <0.0001 | 1.7837 | 0.3049 |
| $F_{SC2}$ | 0.0268 | <0.0001 | 1.2320 | 0.1936 | 0.0323 | <0.0001 | 3.9708 | 0.5911 |
| $F_{ST}$ | 0.0620 | <0.0001 | 1.2835 | 0.1751 | 0.0600 | <0.0001 | 6.1550 | 1.0658 |
| $F_{C1C2}$ | 0.0080 | <0.0001 | 0.1584 | 0.0812 | 0.0130 | <0.0001 | 2.1884 | 0.0948 |
| $F_{C1T}$ | 0.0438 | <0.0001 | 0.2100 | 0.0516 | 0.0412 | <0.0001 | 4.3740 | 0.3788 |
| $F_{C2T}$ | 0.0362 | <0.0001 | 0.0513 | 0.0156 | 0.0286 | <0.0001 | 2.1870 | 0.0947 |

According to Tables 2 and 3, the two kinds of results obtained by using these two methods are generally similar. The variance components within individuals contribute to the majority in these two kinds of results and the *F*-statistics are generally small (below 0.08), implying a medium difference among populations (*F*_*ST*_ ≈ 0.06). For the anisoploid method, the value of inbreeding coefficients is small (*F*_*IS*_ = 0.0119), but it is significantly greater than zero, while it is exactly equal to zero for the likelihood method.

## Discussion

### RMSE

In this paper, we generalize the framework of AMOVA and propose four methods to solve the variance components and the *F*-statistics.

It can be found from the comparison of Figures 2 and 4 that the RMSE in diploids is smaller than that in tetraploids, implying that the estimations of variance components and *F*-statistics are more accurate for diploids, although there are some biases for the polyploid phenotypic data.

We also see from Figures 2 and 4 that for the three moment methods, the estimated *F*-statistic 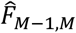 becomes increasingly inaccurate as *M* increases. However, for the likelihood method, as *M* increases, the accuracy of 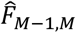 is not only unaffected but also the same as that of 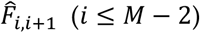. Therefore, the likelihood method can be used in the datasets with higher value of *M*.

### Biasedness

For the homoploid method, the estimated *F*-statistic *F*_*i*,*i*+1_ is biased for the genotypic dataset with missing data. This bias is caused by the weighting for missing data. The allele frequency of the missing genotypes refers to the allele frequency in the corresponding population, and we assume that *F*_*IS*_ = 0. Therefore, *F*_1,2_ is underestimated in Figure 1. For the anisoploid and the weighting genotypic methods, because the missing genotype data are ignored, such a bias is avoided.

For the phenotypic data, all three moment methods become biased. There are two sources of these biases: (i) the extraction of dummy haplotypes; (ii) the estimation of allele frequencies.

The extraction of dummy haplotypes breaks the correlation between alleles within the same individual, which can bias the estimation of SS_WI_. Therefore, the bias of *F*_1,2_ in Figure 3 is largest. This bias can be reduced by increasing the sample size *N* (e.g., the bias is −0.1 at *N* increasing to 50, see Figure 3). For the weighting genotypic method, this bias can be eliminated.

For the genotypic data, the allele frequencies are estimated by counting the alleles within the corresponding genotypes, so this estimation is unbiased. For the phenotypic data, the allele frequencies are estimated by using an expectation-maximization algorithm modified from Kalinowski and Taper (2006). Because this algorithm is also a kind of maximum-likelihood method, such estimation is biased, and the bias is passed to the subsequent steps. However, it can be reduced to a negligible level if *N* is large enough (e.g., the bias is 0.003 at *N* = 50, see Figure 3).

### Unbiasedness

Due to the unbiasedness of moment methods, some negative estimates of variance components and *F*-statistics may present when the level of differentiation is low or the sample size is small. We select three datasets to illustrate this phenomenon, where each dataset consists of two populations with identical diploids which are genotyped at only one biallelic locus. Specifically, in Dataset 1, each population contains four genotypes (1 *AA*, 1 *BB* and 2 *AB*), which are drawn from HWE; in Dataset 2, each population is heterozygote-deficient, only containing two homozygotes (1 *AA* and 1 *BB*); in Dataset 3, each population is heterozygote-excessive, not containing homozygotes (2 *AB*). Because there is only one locus and no missing data, the three moment methods are equivalent. We use the homoploid method as an example, whose results with 9999 permutations are shown in Table 4. The results by using the likelihood method are also shown in this table as a comparison.

**Table 4.**
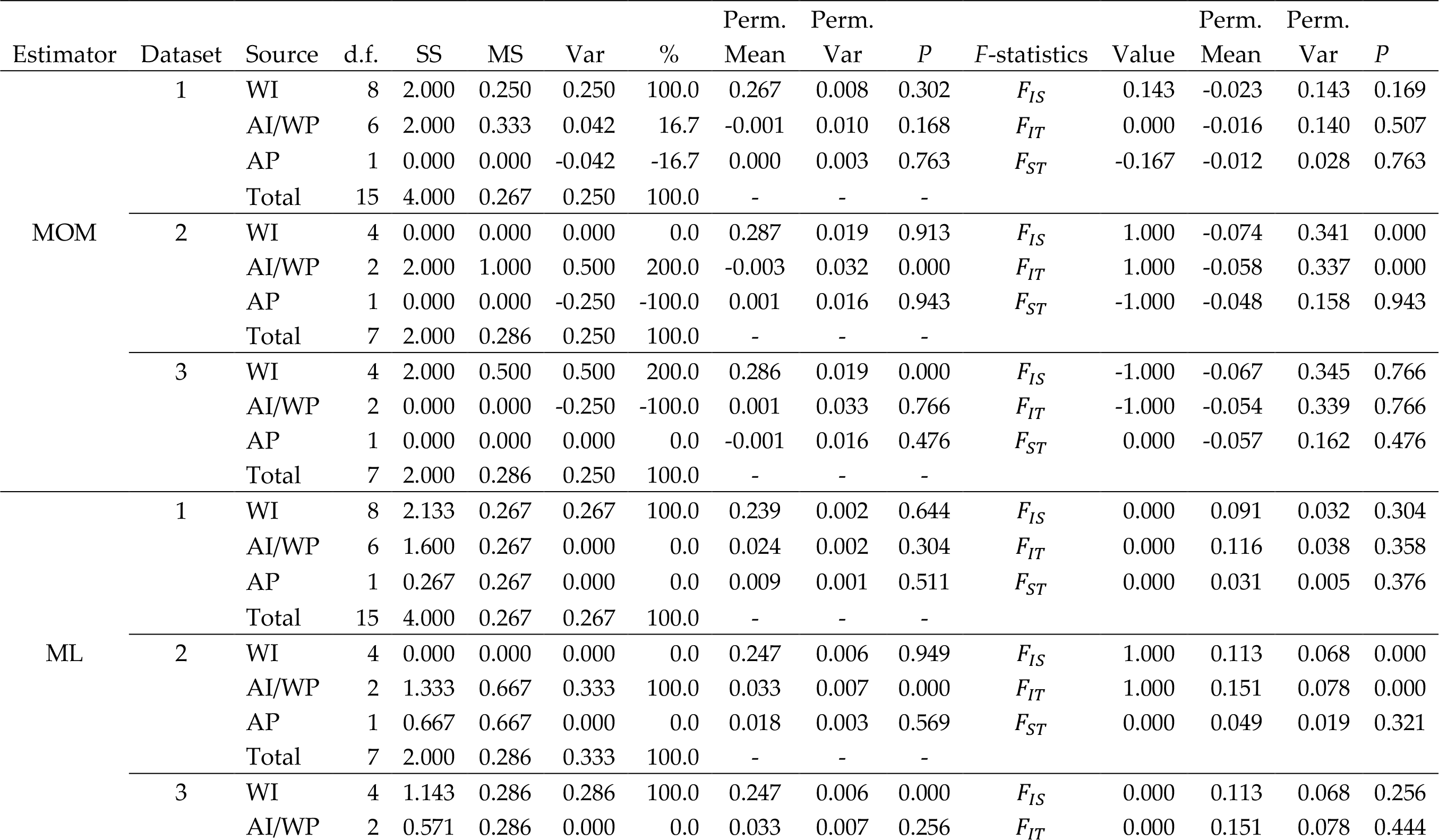
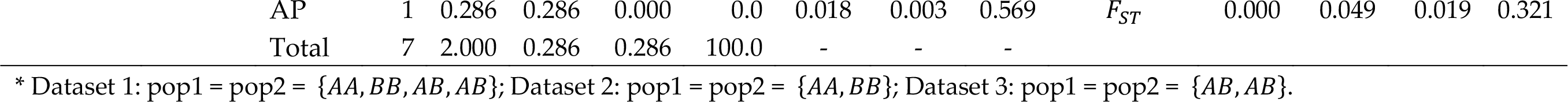
Results of AMOVA by using the moment and likelihood methods

It can be found from Table 4 that for the moment methods, some estimates of variance components and *F*-statistics are negative or greatly deviate from the true values. For the likelihood method, such negative values can be avoided, and the values of estimates can be ensured to lie in the range of biological meaning.

### Empirical results

There are some differences in the results of AMOVA on Pemberton *et al.* (2013) dataset between the moment and the likelihood methods (see Tables 2 and 3). For example, the value of *F*_*IS*_ is significantly positive for the moment method, but it is exactly equal to zero for the likelihood method.

The differences come from the dissimilarity between the schemes of these two kinds of methods. For the moment method, the SS within each hierarchy and at each locus is calculated, and the occurrence of some rare genotypes/phenotypes can only slightly change the values of SS. Therefore, for the loci with a similar polymorphism, their influences on the values of SS, MS, Var etc. are also similar.

In contrast, for the likelihood method, the SS is more sensitive to the distribution of genotypes/phenotypes, and the occurrence of some rare genotypes/phenotypes (e.g., homozygotes of rare alleles) can greatly affect the values of SS. The influence of a single rare genotype/phenotype may be equal to those of thousands of common genotypes/phenotypes. For the loci with a similar polymorphism, their influences on the values of SS, MS, Var etc. may be dramatic.

### Applications

The calculating speed of the homoploid method is fastest during the permutation test. For this method, the genetic distance matrix only needs to be calculated one time, and it is permuted in a very fast way during the permutation test. More specifically, a permutation *k*_1_*k*_2_ … *k*_*H*_ of the number codes 1, 2, …, *H* is randomly generated, where *H* is the order of the genetic distance matrix (i.e., the number of alleles). Let 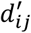 be equal to 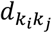, where 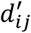 is the *ij*^th^ element in the permuted genetic distance matrix, and 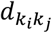 is the 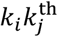 element in the original distance matrix, *i*, *j* = 1, 2, …, *H*. This technique can largely reduce the time expense, especially for a large dataset. For the other methods, the genetic distance should be calculated at each locus and in each iteration, so the calculating speeds of these methods are far slower than that of homoploid method. The drawback of the homoploid method is that the genetic distances are biased for the genotypic dataset with missing data or for the polyploid phenotypic data. Therefore, the homoploid method is suitable for a high-quality genotypic data (with a high genotyping rate) or a large dataset (e.g., next-generation sequencing data).

Although the calculating speed of the anisoploid method is slower than that of the homoploid method, it is still faster than those of the other two methods. That is because the whole dataset needs to be regenerated during the permutation test in the other two methods. For the anisoploid method, because the missing data are ignored during the calculation, the genetic distances are unbiased for genotypic data with a low genotyping rate. Therefore, this method is suitable for a low-quality genotypic data.

For the weighting genotypic method, because no dummy haplotypes are extracted, the genetic distances are less biased for the phenotypic data. In this method, instead of the use of permutation test, it randomly generates the dataset under the hypothesis that there is no differentiation. After that, it also calculates the probability that the variance components or the *F*-statistics at each locus and in each iteration are greater than the observed values. Therefore, this method is suitable for the polyploid phenotypic data.

For the three moment methods, there are two problems: (i) the RMSE of each 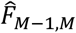 increases as the hierarchy number *M* increases, and (ii) some negative variance components or some negative estimates of *F*-statistics may present when the difference due to the unbiasedness is small. For the likelihood method, these two problems can be avoided, and the RMSEs of the estimated *F*-statistics are insensitive to *M*. In addition, various values of estimation are always in the biologically meaningful range. Therefore, this method is suitable for a larger *M* and/or for datasets for which a part of the results obtained by using these moment methods cannot be explained.

## Acknowledgment

KH would like to thank Prof. Kermit Ritland for providing the space of visiting scholar in the University of British Columbia. This study was funded by the Strategic Priority Research Program of the Chinese Academy of Sciences (XDB31020302), the National Natural Science Foundation of China (31770411, 31730104, 31572278 and 31770425), the Young Elite Scientists Sponsorship Program by CAST (2017QNRC001), and the National Key Programme of Research and Development, Ministry of Science and Technology (2016YFC0503200). DWD is supported by a Shaanxi Province Talents 100 Fellowship.

## Author contributions

KH and BGL designed the project, KH and YLL constructed the model and wrote the draft, KH designed the software, PZ performed the simulations and analyses, and DWD checked the model and helped to write the manuscript.

## Supplementary materials

### Linear model

Let *A* be an allele randomly taken from the total population, and let *p* be the mean frequency of *A* in the total population. We will focus on the biases of the frequency *p* related with an allele *a* to carry out our discussion. Our linear model is developed from Cockerham (1969; 1973), which is described by the following function:

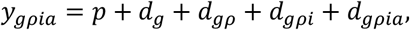

where *a* is arbitrary, and the relations among *g*, *ρ*, *i*, *a* are nested, that is, *a* ∈ *i* ⊆ *ρ* ⊆ *g*; *d*_*g*_ is the bias of the frequency *p* in the group *g* relative to the total population, *d*_*gρ*_ is the bias of *p* in the population *ρ* relative to the group *g*, *d*_*gρi*_ is the bias of *p* in the individual *i* relative to the population *ρ*, and *d*_*gρia*_ is the bias of *p* in the allele *a* relative to the individual *i*. It is worth pointing out that because of the nested relation, *g* and *ρ* are uniquely determined so long as *i* is given. We stipulate that the condition E(y_*gρia*_) = *p* should be satisfied in this model.

Because E(y_*gρia*_) = *p* and the allele frequencies obey a binomial distribution, we have var(*y*_*gρia*_) = *p*(1 − *p*), that is, 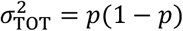.

According to Cockerham (1969; 1973), the formulas for cov(*y*_*gρia*_, *y*_*g′ρ′i′a′*_) under various situations are

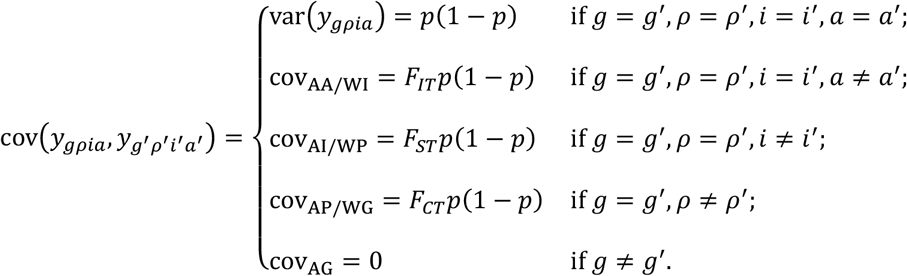

For the final situation, because the alleles among groups are assumed to be independent, the value of corresponding *F*-statistic is zero, and so cov_AG_ = 0*p*(1 − *p*) = 0. Moreover, the formulas for E(*y*_*gρia*_*y*_*g′ρ′i′a′*_) under various situations are

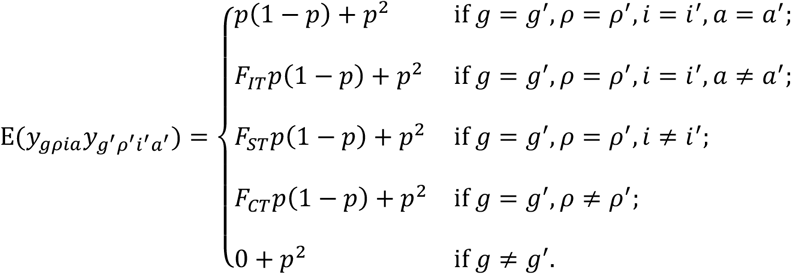

In fact, for the first situation, E(*y*_*gρia*_*y*_*g′ρ′i′a′*_) becomes 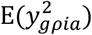, so

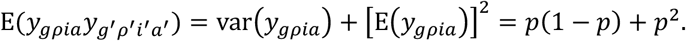

For the second situation, E(*y*_*gρia*_*y*_*g′ρ′i′a′*_) becomes E(*y*_*gρia*_*y*_*gρia′*_). Because *F*_*IT*_ is the probability Pr(*y*_*gρia*_ ≡ *y*_*gρia′*_) in the sense that two distinct alleles within a same individual in the total population are IBD, we obtain

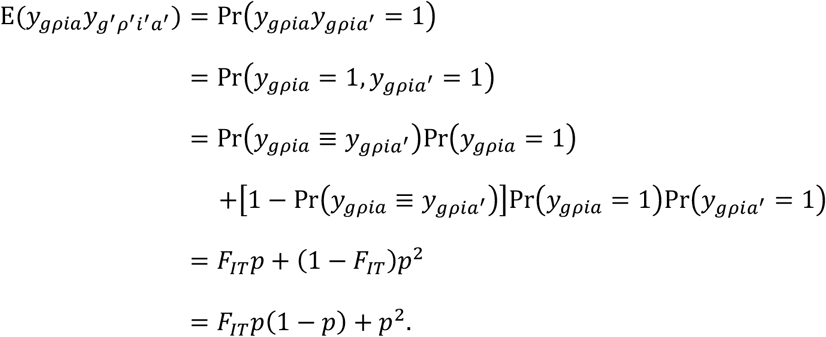

For the remaining situations, the derivations are similar, and omitted.

There are the following relations between the *F*-statistics and the variance components:

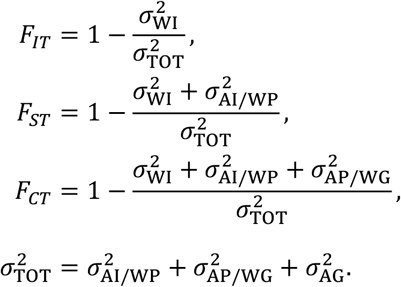

Because 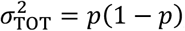, we obtain

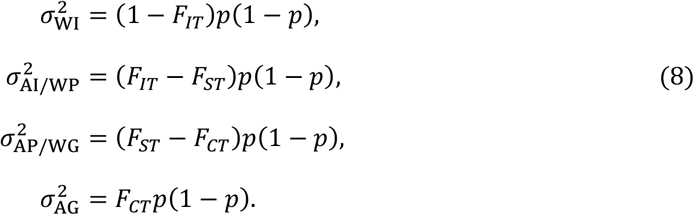

We will use the symbol 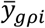 (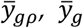 or 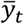) to denote the average of values of the function *y*_*gρia*_ when *a* is taken from all alleles within the individual *i* (the population *ρ*, the group *g* or the total population). Then,

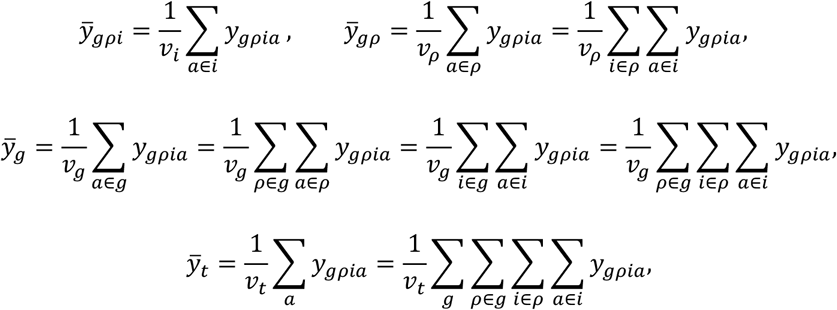

where *ν*_*i*_, *ν*_*g*_, *ν*_*ρ*_ or *ν*_*t*_ is the number of alleles within the individual *i*, the population *ρ*, the group *g* or the total population, respectively.

### Derivation for the formula of expected SS_WI_

The expectations of 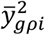 and 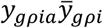 are calculated as follows:

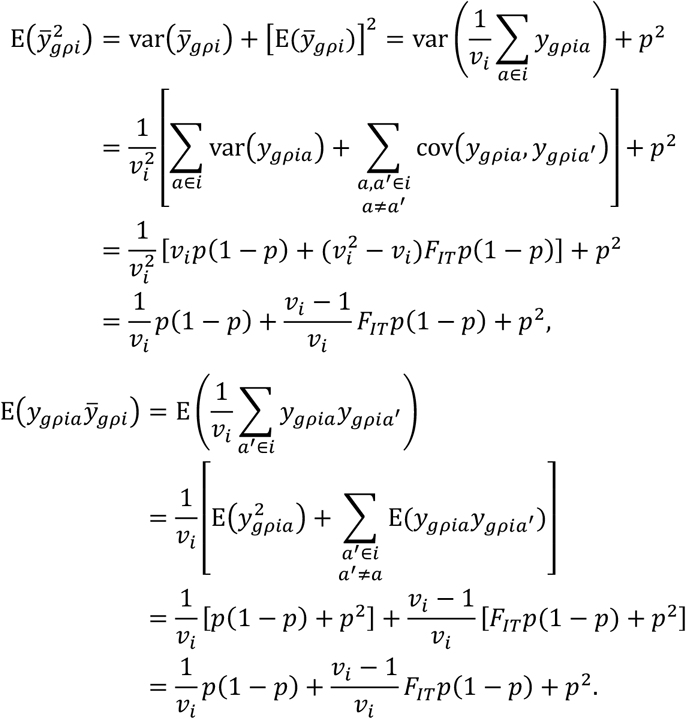

Comparing the two calculated results, we have 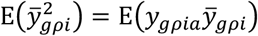. In addition,

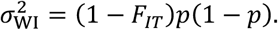

Using these facts, we can easily obtain the next result:

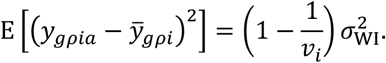

Moreover, we have

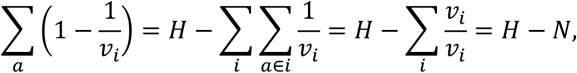

where *H* and *N* are the numbers of alleles and individuals in the total population, respectively. Now, we can easily derive the first formula of expected SS in Equation (2):

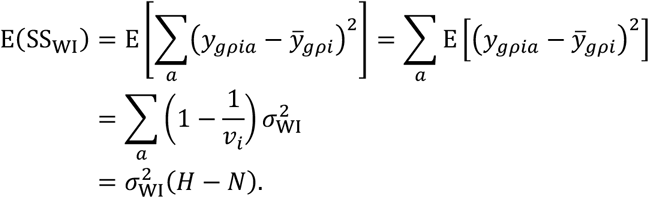

### Derivation for the formula of expected *SS*_WP_

The expectations of 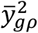 and 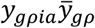 are calculated as follows:

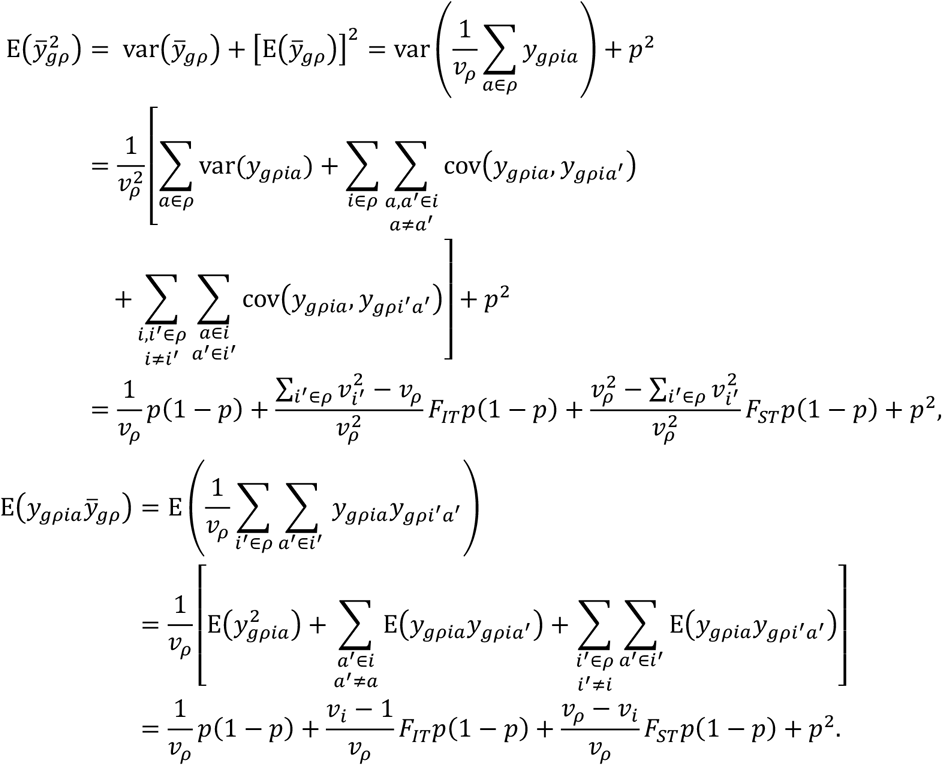

Now, by using Equation (8), it is not difficult to calculate that

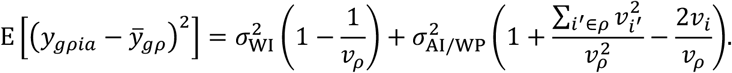

Moreover, because *P* is the number of populations in the total population, we have

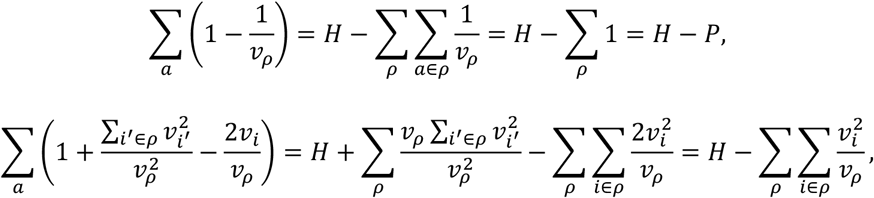

Now, the second formula of expected SS in Equation (2) can be derived as follows:

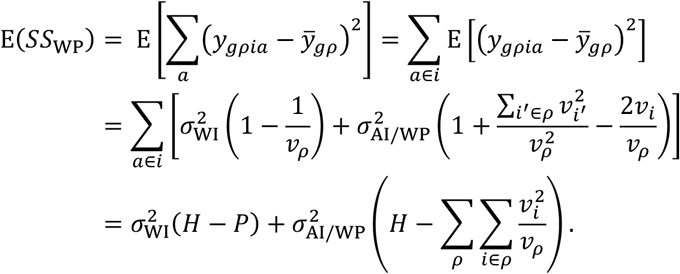

### Derivation for the formula of expected *SS*_WG_

The expectations of 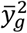 and 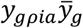 are calculated as follows:

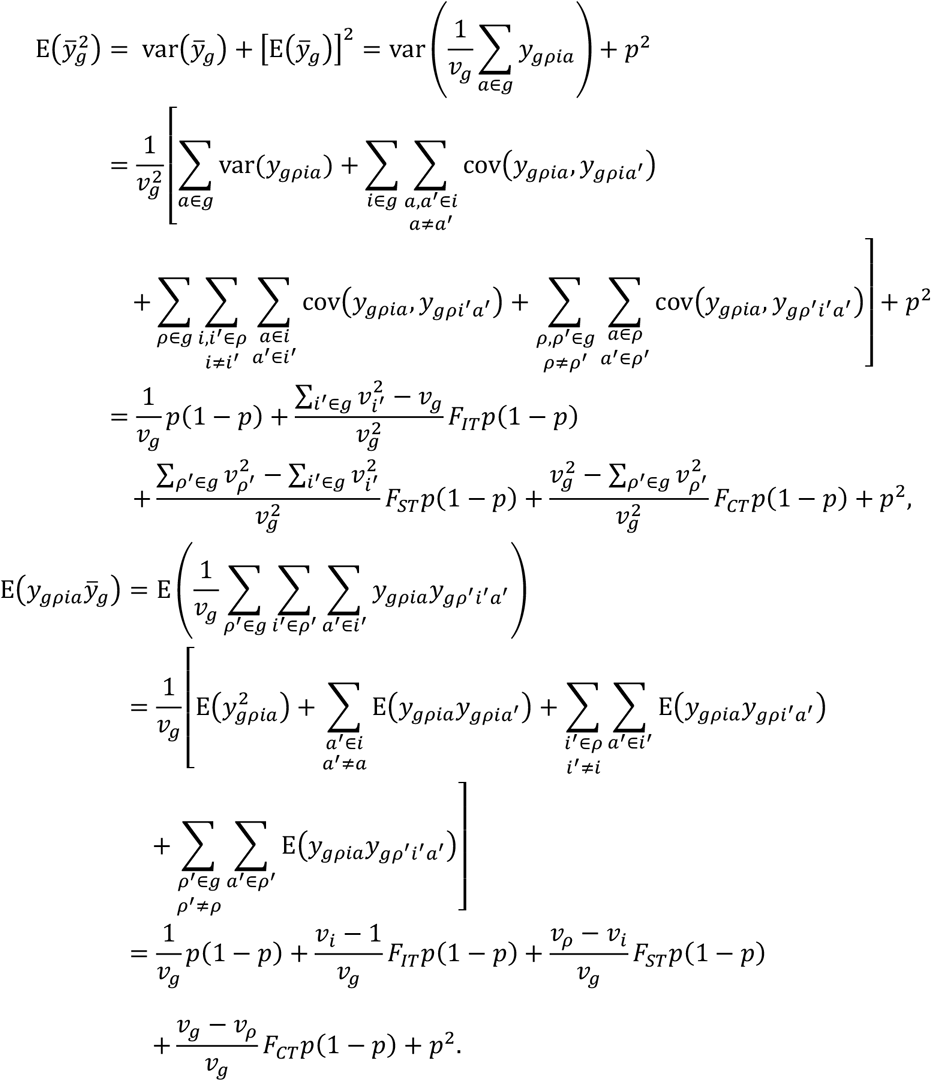

Now, by using Equation (8), it is easy to calculate that

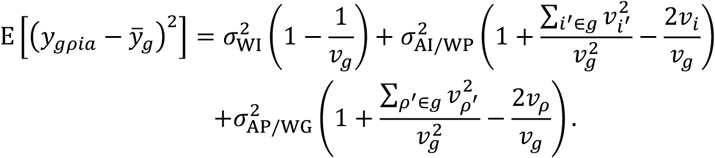

Moreover, because *G* is the number of all groups, if we let *x* = *i* or *x* = *ρ*, then

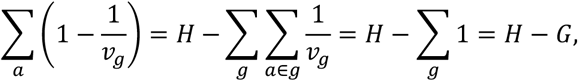

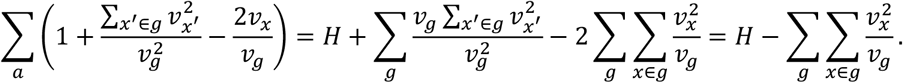

Now, the third formula of expected SS in Equation (2) can be derived as follows:

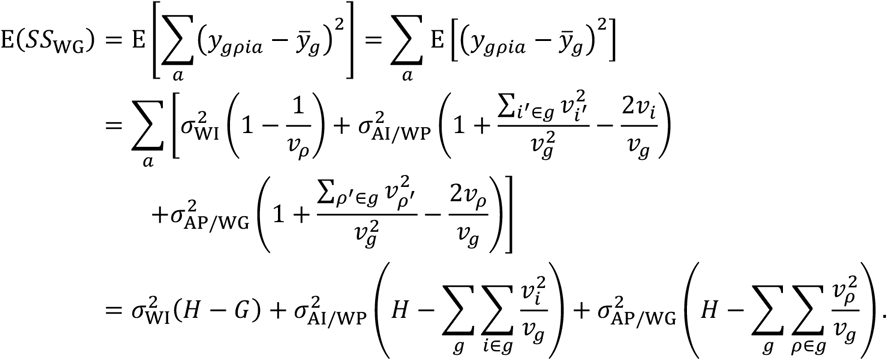

### Derivation for the formula of expected *SS*_TOT_

The expectations of 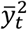 and 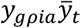 are calculated as follows:

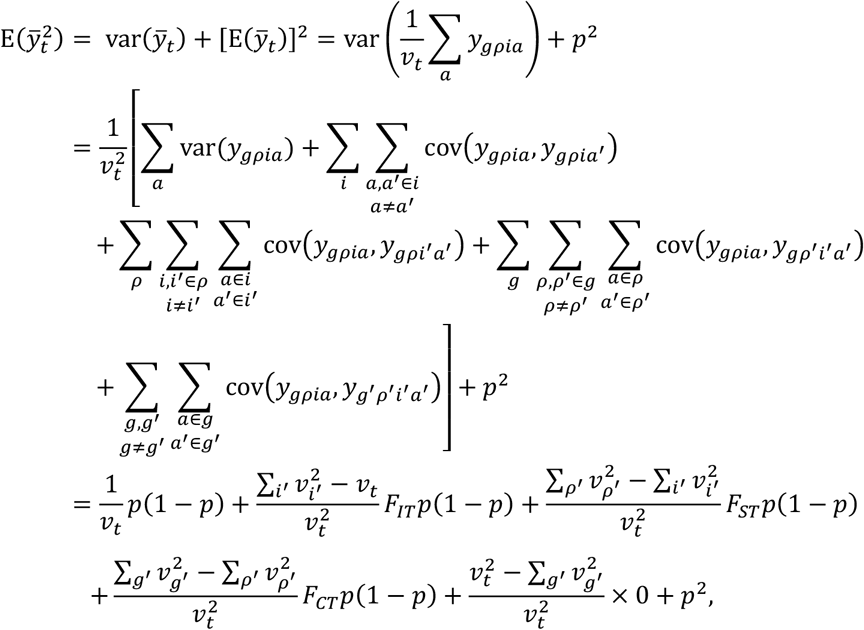

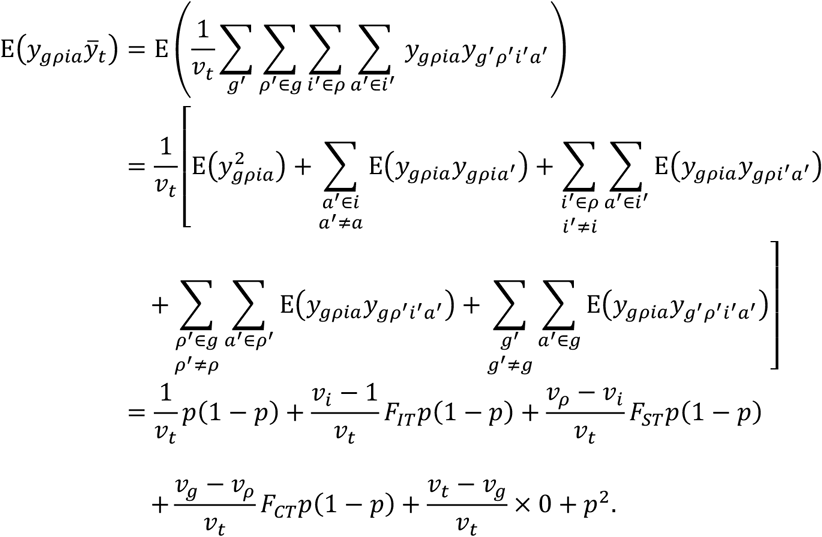

Now, according to Equation (8), we can easily calculate that

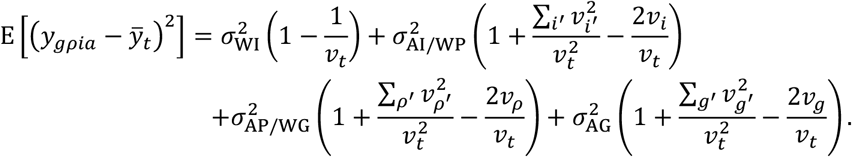

Moreover, we have 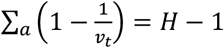 and

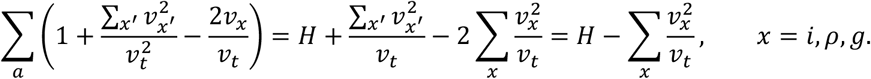

Now, let us derive the final formula in Equation (2):

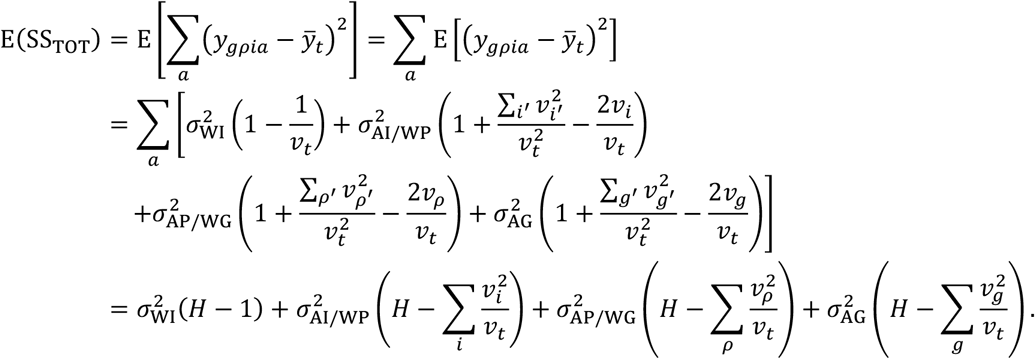

